# Local adaptation of host-species specific gut microbiota

**DOI:** 10.1101/2022.08.22.504808

**Authors:** Daniel D. Sprockett, Jeffrey D. Price, Anthony F. Juritsch, Robert J. Schmaltz, Madalena V. F. Real, Samantha Goldman, Michael Sheehan, Amanda E. Ramer-Tait, Andrew H. Moeller

## Abstract

Mammalian species harbor compositionally distinct gut microbial communities, but the mechanisms that maintain specificity of symbionts to host species remain unclear. Here we show that natural selection within house mice (*Mus musculus domesticus*) drives deterministic assembly of the house-mouse gut microbiota from mixtures of native and non-native microbiotas. Competing microbiotas from wild-derived lines of house mice and other mouse species (*Mus* and *Peromyscus* spp.) within germ-free wildtype (WT) and *Rag1*-knockout (*Rag1*^−/−^) house mice revealed widespread fitness advantages for native gut bacteria. Certain native Bacteriodetes and Firmicutes favored by selection in WT hosts were not favored or disfavored in *Rag1*^−/−^ hosts, which lack adaptive immunity, indicating that *Rag1* mediates fitness advantages of these strains. This study demonstrates local adaptation of gut microbiota to a mammalian species.

**One-Sentence Summary:** Adaptive advantages for native bacteria underlie the assembly of the mouse gut microbiota.

## Main Text

The gut microbial communities of diverse mammalian species reflect the evolutionary histories of their hosts. In rodents, xenarthra, artiodactyls, primates, and other mammalian clades, hosts of the same species (*i.e*., conspecifics) tend to harbor more similar gut microbiota compositions than do hosts of different species, and microbiota dissimilarity between host species is positively associated with the hosts’ evolutionary divergence (*1–7*). Experimental work in rodents has indicated that disruption of host-species specific microbiotas can have adverse consequences for hosts, including impaired resistance to pathogen colonization (*8*), diminished nutrient utilization (*9*), and reduced growth rate (*10*). Nevertheless, the ecological and evolutionary forces underlying microbiota specificity to host species have not been determined (*11–14*). One proposed mechanism is biased microbial dispersal (*i.e*., dispersal limitation). Microbial dispersal among conspecific hosts occurs readily through both social (*15–18*) and vertical transmission (*19–23*), whereas dispersal between host species tends to be less frequent (*3, 24*). The bias towards dispersal among conspecific hosts could promote and maintain microbiota divergence between host species in the absence of selective processes (*3, 13*). A non-mutually exclusive mechanism is adaptation of symbionts to their respective host species (*i.e*., local adaptation). For example, previous experiments have shown that strains of the gut bacterium *Lactobacillus reuteri* derived from house mice (*Mus musculus domesticus*) display higher fitness within house mice than do *L. reuteri* strains from other mammalian host species (*25–27*). However, the extent to which constituents of host-species specific microbiotas are favored by natural selection within their host species remains unclear. Quantifying the relative influences of these alternative mechanisms—dispersal limitation and local adaptation—remains a critical gap in understanding the assembly of host-species specific microbiota in mammals.

We tested for local adaptation of the house-mouse gut microbiota through a series of *in vivo* microbiota competition experiments. First, we characterized the gut microbiotas of nine laboratory mouse lineages of house mice and other *non-domesticus* mouse species. These included the house mouse model C57BL/6 and eight wild-derived lineages, which represented three species within the genus *Mus* (*M. m. domesticus, M. spicilegus*, and *M. pahari*) and one species of deer mouse (*Peromysucs maniculatus*) (Table S1) (Fig. 1A). Mouse lineages were maintained under laboratory conditions for >10 host generations. Wild-derived lineages were descended directly from mice collected in the wild (*i.e*., they were never rederived through embryo transplantation to or cross-fostering with a laboratory mouse line) to facilitate the retention of host-lineage specific microbiota (*10, 23*). Amplicon sequencing of the 16S rRNA gene from fecal samples indicated that host species maintained compositionally distinct microbiotas in the lab environment (Fig. S1A and B, PERMANOVA p <0.001). Furthermore, microbiota similarity between hosts decayed exponentially as a function of host evolutionary divergence time (Fig. 1B; R^2^ = 0.54, p <0.001), recapitulating what has been observed in wild rodent species (*3*). Mean microbiota similarity between individual hosts was highest within host strains, followed by between strains/within species, between species, and between genera (Fig. 1B, inset). A significant negative relationship between microbiota similarity and host relatedness was also observed within the genus *Mus* alone (Fig. S2; R^2^ = 0.19, p <0.001). Together, these results indicate that the laboratory mouse lineages have maintained host-species specific microbiotas associated with their hosts’ phylogenetic histories.

**Fig. 1.**
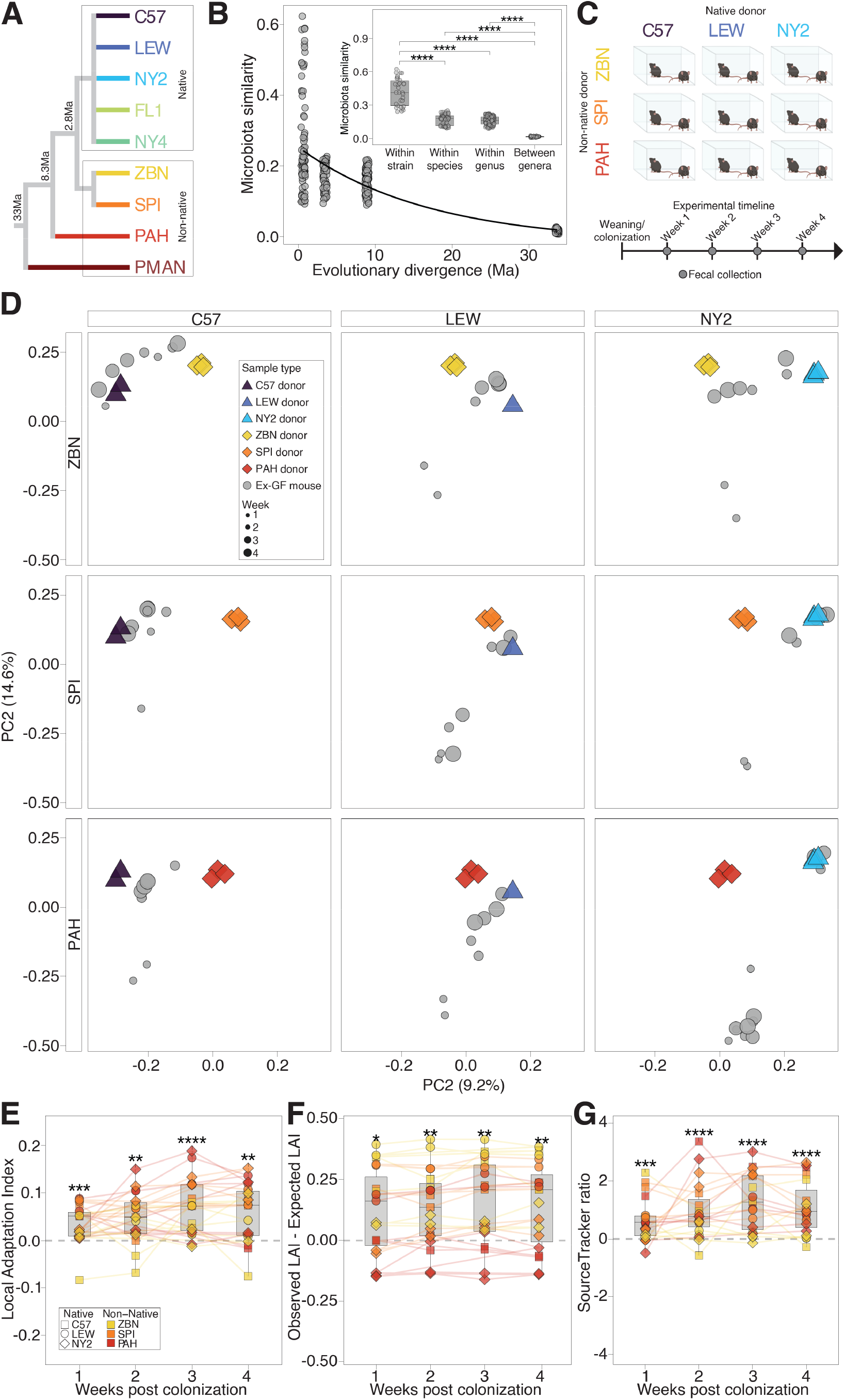
Deterministic assembly of house-mouse microbiota from mixtures of native and non-native microbiotas. (**A**) Phylogeny shows evolutionary relationships among wild-derived lab mice from which microbiotas were obtained. (**B**) Scatterplot shows negative association between microbiota similarity (Jaccard) and evolutionary divergence (Ma, millions of years ago) among rodent lineages. Dashed line shows exponential decay regression (p <0.001, R^2^ = 0.54). Inset shows boxplots of microbiota similarity between pairs of samples. Wilcoxon Tests, FDR-adjusted p-values **** <1 × 10^−4^. (**C**) Cartoon shows experimental design. Fecal microbiotas from three native *Mus musculus domesticus* lines and three non-*domesticus* mouse lineages were mixed in pairwise combinations and inoculated into weaned germ-free mice, from which fecal samples were collected weekly for 4 weeks. (**D**) Principal Coordinate Analysis plots show similarities among microbiotas from donors and ex-germ-free recipients based on the Jaccard similarity index. Colors indicate the mouse lineage from which the samples were collected corresponding to (**A**). Sizes of grey circles indicate time points (weeks 1 through 4). Facets plotted along common axes show microbiotas from individual pairs of donors and their corresponding recipient ex-germ-free mice. (**E**) Boxplots show positive Local Adaptation Index (LAI) values of ex-germ-free mice throughout the 4-week experiment; (**F**) positive differences between observed LAI values and LAI values expected under neutral assembly; and (**G**) log_10_ transformed ratios of native ASVs to non-native ASVs identified as sources by SourceTracker. In (**E**–**G**), shapes and colors denote identities of native and non-native donors, respectively, and lines connect samples from the same mouse. For each boxplot in (**B, E**–**G**), center lines denote medians, and lower and upper hinges correspond to first and third quartiles, respectively. FDR-adjusted *p*-values were derived from Wilcoxon tests for non-zero mean, * <0.05, ** <0.01, *** <0.001, **** <0.0001.

Using samples from the diverse set of mouse lineages, we then conducted competition experiments in which microbiotas from house mice (*i.e*., native microbiota) and from non-*domesticus* mouse species (*i.e*., non-native microbiota) were co-inoculated at defined ratios into germ-free house mice. The first set of experiments tested nine distinct pairwise mixtures of native and non-native microbiotas derived from three of the *M. m. domesticus* mouse lineages (*i.e*., native donors: C57BL/6, LEWES, and NY2) and three *non-domesticus* mouse lineages (*i.e*., non-native donors: *M. spicilegus* ZBN and SPI/TUA, and *M. pahari* PAH) (Fig. 1C). Pairs of native and non-native fecal samples were mixed equally by weight, and each mixture was inoculated at weaning into two germ-free *M. m. domesticus* (C57BL/6) hosts reared together in an individual microisolator cage (Fig. S3A). Fecal samples from ex-germ-free recipient mice were collected weekly for four weeks and profiled using 16S rRNA gene amplicon sequencing. Microbiota similarity (Jaccard) between ex-germ-free mice and their corresponding donor fecal samples was significantly higher on average than was that between the ex-germ-free mice and fecal samples from other donors (Fig. 1D, Fig. S4; Wilcoxon Test, FDR adjusted p-value <1 × 10^−5^), indicating successful inoculations.

To test for local adaptation, we calculated microbiota similarity between ex-germ-free mice and their native or non-native donors. We defined the difference between the microbiota similarity to the native donor and the microbiota similarity to the non-native donor as the Local Adaptation Index (LAI) (Fig. S3B; Supplementary Materials). LAI values were significantly positive for ex-germ-free mouse microbiotas at every time point (Fig. 1E, Table S2), indicating that microbiota similarity to native donors was higher than that to non-native donors. The increased microbiota similarity to native donors was also evident in a Principal Coordinates Analysis (PCoA) (Fig. 1D). Positive LAI values of ex-germ-free mouse microbiotas are consistent with local adaptation of native microbiotas relative to non-native microbiotas. However, differences in bacterial load between donor fecal samples could lead to differences in colonization success between native and non-native microbiotas in the absence of local adaptation. Moreover, positive LAI could result from differences in alpha diversity between the native and non-native microbiotas included in the mixtures, even if the microbiotas colonized ex-germ-free house mice equally well. To address these potential issues, we calculated the expected LAI for each ex-germ-free mouse microbiota under a neutral model of community assembly (*28*) given the measured microbiota composition and bacterial load (*i.e*., 16S rRNA gene copy number) of the two donor fecal samples included in the mixture that the mouse received (Table S3; Supplementary Materials). The difference between the observed LAI and the expected LAI provided a test statistic for the degree of competitive advantage of the native microbiota over the non-native microbiota (Fig. S3C; Table S2). Results showed that native microbiotas were selected over non-native microbiotas in ex-germ-free house mice throughout the four week experiments beyond what would be expected under neutrality (Fig. 1F; Wilcoxon Tests, FDR adjusted p-value <0.05 for all comparisons; Observed LAI – Expected LAI > 0). In addition, parallel analyses based on SourceTracker (*29*) indicated that ex-germ-free mouse microbiotas were composed of a significantly greater microbiota fraction derived from the native donor than from the non-native donor (Fig. 1G; Wilcoxon Tests, FDR adjusted p-value <0.001 for all comparisons) (Supplementary Materials). Cumulatively, these results show that, when competed within germ-free house mice, native house-mouse microbiotas significantly outcompeted nonnative microbiotas from *non-domesticus Mus* host species.

Next, we tested whether competitive advantages for native microbiotas over non-native microbiotas depend on the presence of the host adaptive immune system. Adaptive immunity in mammals is both a highly specific microbial filter as well as a facilitator of colonization for certain microbiota constituents (*30, 31*). To assess the degree to which host adaptive immunity contributes to selection for host-species specific microbiota, we conducted additional microbiota competition experiments in which two mixtures of native and non-native microbiotas—*M. m. domesticus* (FL1) +*M. pahari* (PAH) and *M. m. domesticus* (NY4) + *Peromyscus maniculatus* (PMAN)—were inoculated into wildtype (WT) germ-free C57BL/6 mice (n=34) and a line of germ-free C57BL/6 in which *Rag1* gene has been deleted (*Rag1*^−/−^) (n=40) (Fig. 2A). *Rag1*^−/−^ mice lack mature T and B cells and a functional adaptive immune system (*32*), and deletion of *Rag1* has previously been shown to alter the microbiota relative to WT hosts (*30, 31, 33, 34*). Here, each microbiota mixture was inoculated into a single, sterile isolator containing 5-7 cages and 2-4 mice per cage. Mice of the same sex and genotype were cohoused, and cages were arranged in a block design with respect to genotype within isolators. Fecal samples from ex-germ-free mice were collected at four- and six-weeks post-inoculation, and microbiota profiles were generated with 16S rRNA gene sequencing. As in the first set of experiments, microbiota similarity (Jaccard) between the ex-germ-free mice and the donor fecal samples that they received was significantly higher on average than that between the ex-germ-free mice and other donor fecal samples (Fig. 2B, Fig. S5; Wilcoxon Test, FDR adjusted p-value <1 × 10^−5^).

**Fig. 2.**
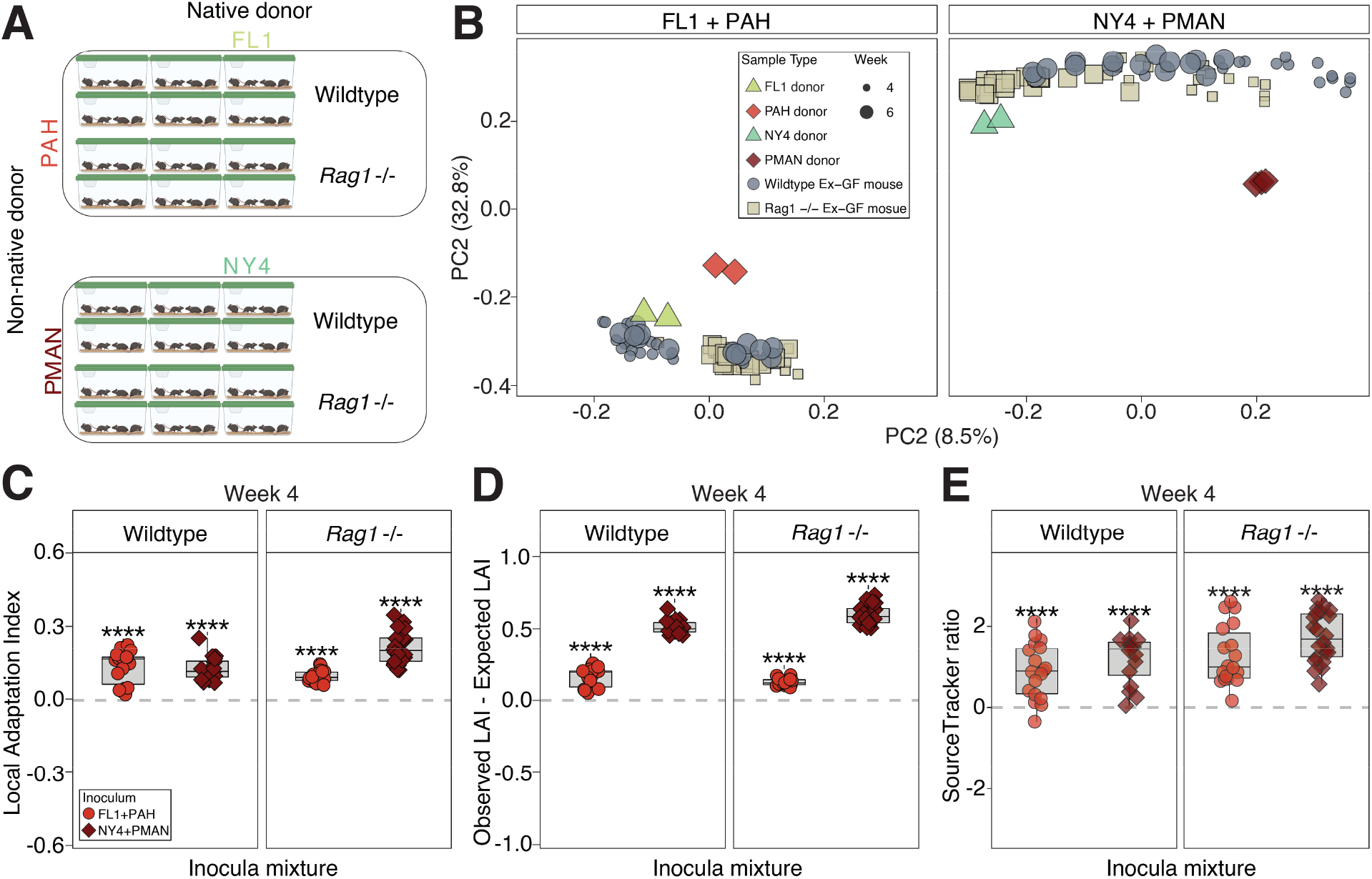
Competitive advantages of native microbiota in both WT and *Rag1*^−/−^ mice. (**A**) Cartoons show experimental design. Fecal samples from two *Mus musculus domesticus* lines and two non-*domesticus* mouse strains were mixed in equal ratios and inoculated into germ-free mouse pups at 10 days of age reared in sterile, multi-cage gnotobiotic isolators. Fecal samples were collected at 4 and 6 weeks. (**B**) Principal Coordinate Analysis plots show microbiota similarities (Jaccard) among donors and ex-germ-free recipients. Colors indicate mouse lineages from which samples were collected. Sizes of grey circles indicate time points (weeks 4 and 6). Facets plotted along common axes show microbiotas from individual pairs of donors and their recipient ex-germ-free mice. (**C–E**) Boxplots show Local Adaptation Index (LAI) values (**C**), the differences between observed and expected LAI values (**D**), and log_10_ transformed ratios of native ASVs to non-native ASVs identified as sources by SourceTracker (**E**). For (**C**–**E**), shapes and colors denote identities of native and non-native donors, respectively. Wilcoxon test for nonzero mean, FDR-adjusted p-values * <0.05, ** <0.01, *** <0.001, **** <0.0001. For each boxplot in (**B**–**D**), center lines denote medians, and lower and upper hinges correspond to first and third quartiles, respectively.

This second set of experiments revealed reproducible assembly of the house-mouse gut microbiota from mixtures of native and non-native microbiotas regardless of the presence or absence of a functional adaptive immune system in hosts. Alpha- and beta-diversity differed between ex-germ-free WT and *Rag1*^−/−^ mice (Fig. S6 Wilcoxon Tests, p <0.05 for both inocula; Fig. 2B, PERMANOVA, p <0.001), but significant evidence for local adaptation was observed in both host genotypes. LAI values of ex-germ-free mice were positive at both at four-weeks (Fig. 2C) and six-weeks (Fig. S7A) (Wilcoxon Test, FDR adjusted p-value <0.001 for all comparisons). Moreover, in both experiments and both host genotypes, the observed LAI was significantly greater than expected LAI (Fig. 2D, S7B; Wilcoxon Tests, FDR adjusted p-value <0.001 for all comparisons; Observed LAI – Expected LAI > 0), and microbiotas of ex-germ-free mice contained a greater microbiota fraction originating from the native donor than from the non-native donor (based on SourceTracker) (Fig. 2E, Fig. S7C; Wilcoxon Tests, FDR adjusted p-value <0.001 for all comparisons). These results confirm local adaptation of house-mouse gut microbiota to house mice, and further demonstrate competitive advantages for native microbiota independent of host adaptive immunity.

One possible explanation for the observed competitive advantages of house mouse microbiota over non-native microbiotas is a fundamental inability of non-native microbiotas to colonize germ-free house mice. However, additional experiments in which microbiotas from mouse lineages were inoculated individually into germ-free C57BL/6J revealed no consistent difference in colonization success between native and non-native microbiotas (Supplementary Materials) (Fig. S8). Hundreds of non-native ASVs were detected in individually inoculated ex-germ-free house mice and their corresponding non-*domesticus* donor but not detected in ex-germ-free house mice that received fecal mixtures containing samples from the non-*domesticus* donor (Supplementary Materials). Thus, house mice represent potential niche space (*35*) for many of the gut bacterial lineages from non-*domesticus* mouse species; however, these non-native bacteria tend to be excluded when inoculated in a competitive context with gut bacteria derived from house mice.

Given the widespread evidence of local adaptation of the house-mouse gut microbiota, we next identified the individual gut bacterial lineages that were favored by selection in hosts. We calculated the expected relative abundance in ex-germ-free mice for every amplicon sequence variant (ASV) in the FL1+PAH and NY4+PMAN fecal mixtures (Fig. 2A) under a model of neutral assembly based on the compositional profiles and bacterial loads of the donor samples (Supplementary Materials). Comparing the observed ASV frequencies with the expected ASV frequencies revealed dozens of ASVs that displayed consistent deviations from neutrality in both WT and *Rag1*^−/−^ ex-germ-free mice (binomial tests, *p*-value <0.05) (Table S5). The proportion of ASVs that displayed significant positive deviations from neutrality in competition experiments was significantly higher for native ASVs than for non-native ASVs (Chi-squared test, p-value <0.001), further confirming the local adaptation of native ASVs. Native ASVs favored by selection based on binomial tests (*p*-value <0.05) were significantly overrepresented within Bacteroidetes relative to Firmicutes (Chi-squared *p*-value <1 × 10^−5^) or to all non-Bacteroidetes lineages (Chi-squared *p*-value <1 × 10^−5^) (Fig. 3A). Taxonomic assignments and results of selection analyses for all ASVs are presented in Table S5.

**Fig. 3.**
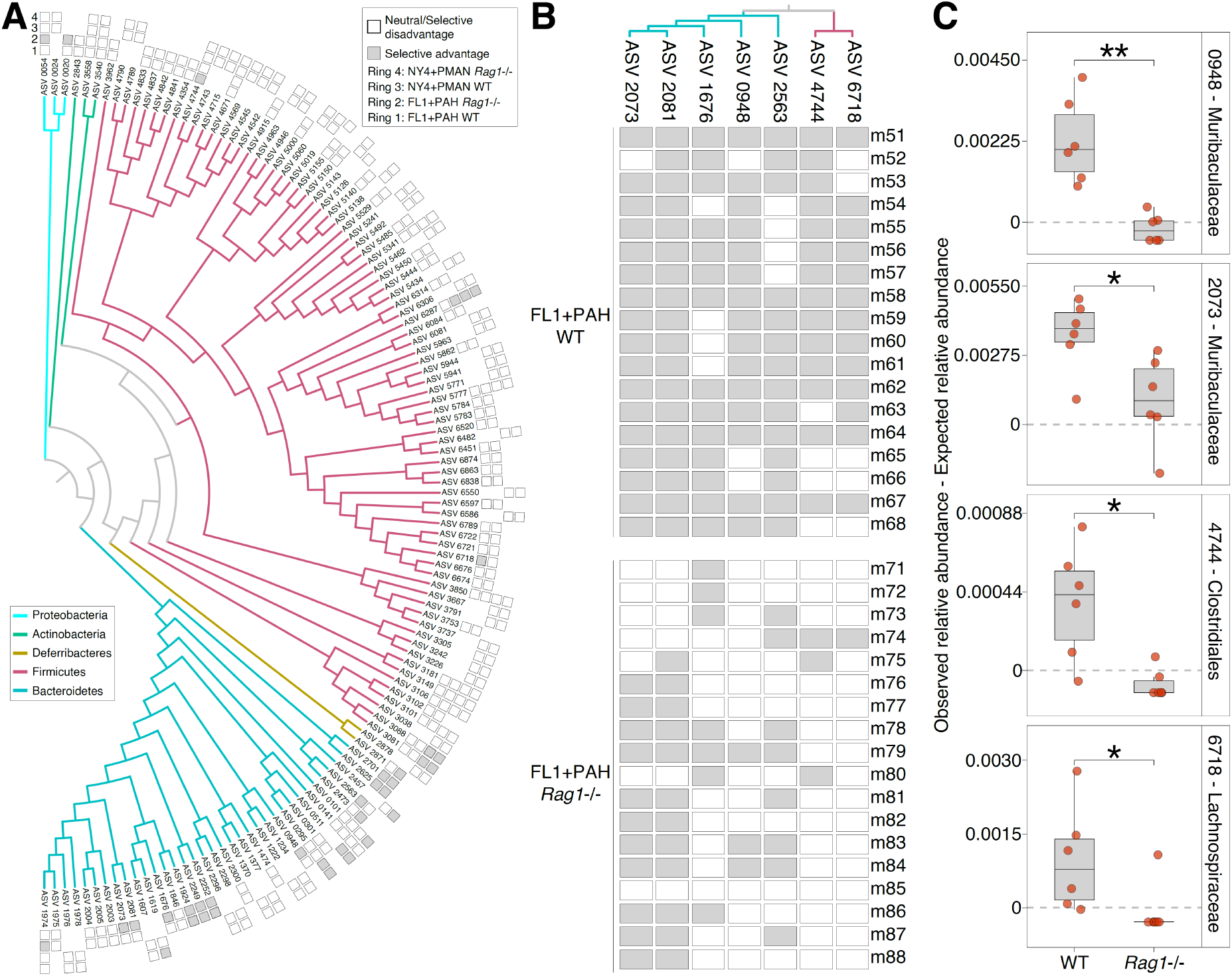
Selective advantages for a subset of native ASVs depended on *Rag1*. (**A**) Phylogeny shows relationships among *Mus musculus domesticus*-specific ASVs detected in ex-germ-free mice that received the NY4+PMAN or FL1+PAH microbiota mixtures. Colors of branches denote bacterial phyla. Rings correspond to ex-germ-free mouse groups (innermost: FL1+PAH WT; second from innermost: FL1+PAH *Rag1*^−/−^ second from outermost: FL1+PAH WT, and outermost: FL1+PAH *Rag1*^−/−^) and indicate significant positive selection on ASVs (filled squares) within ex-germ-free mice based on binomial tests. Unfilled squares mark ASVs that were detected in ex-germ-free mice but not significantly positively selected. (**B**) Phylogeny from (**A**) pruned to only ASVs displaying significant selective advantages in WT ex-germ-free mice but not in *Rag1*^−/−^ mice. Rows correspond to individual ex-germ-free mice. Filled squares indicate ex-germ-free mice in which the observed relative abundance of the ASV exceeded the relative abundance expected under neutrality. (**C**) Boxplots display differences between observed and expected ASV relative abundances in WT and *Rag1*^−/−^ mice that received the FL1+PAH mixture. All ASVs in (**B**) for which cage-mean differences between host genotypes remained significant after false discovery rate (FDR) correction are shown. FDR-adjusted p-values * <0.05, ** <0.01.

Comparing outcomes for native ASVs between WT and *Rag1^−/−^* ex-germ-free mice indicated that, in most cases, selective advantages did not depend on host adaptive immunity (Fig. 3A), consistent with results of beta-diversity-based analyses (Fig. 2). However, seven native ASVs were significantly favored by selection in WT hosts (binomial tests, *p*-value <0.05) but not in *Rag1*^−/−^ hosts (binomial tests, *p*-value > 0.05) (Fig. 3A, B). The negative effects of *Rag1* deletion on ASV-specific selective advantages were only detected within mice that received the FL1+PAH mixture (*i.e*., none were detected within mice that received the NY4+PMAN mixture). Four out of seven of these ASVs displayed significantly greater positive deviations in relative abundance from neutrality in WT mice than in *Rag1*^−/−^ mice based on Wilcoxon tests of per-cage mean relative abundances (Benjamin-Hochberg corrected *p*-values <0.05) (Fig. 3C) (Supplementary Materials). Thus, for a subset of native gut bacterial lineages, selective advantages within house mice depended on the presence of *Rag1*, suggesting selection on native ASVs mediated by the host adaptive immune system.

Microbiota competition experiments in germ-free mice, in which microbial dispersal events were controlled, enabled tests for local adaptation of the house-mouse gut microbiota. In all sets of experiments conducted, native microbiotas consistently outcompeted non-native microbiotas within house mouse hosts. These results demonstrate natural selection favoring the assembly of a host-species specific mammalian gut microbiota.

## Acknowledgments

We thank Dr. Weiwei Yan for assistance with DNA extractions from rodent fecal samples, Dr. Joao Carlos Gomes Neto for assistance with germ-free mouse pup inoculations, and the staff at the Nebraska Gnotobiotic Mouse Program for outstanding animal care.

## Funding

Funding was provided by the National Institutes of Health grant R35 GM138284-01 (AHM).

## Author contributions

Conceptualization: DDS, AR-T, AHM
Methodology: DDS, JDP, AR-T, AHM
Investigation: DDS, JDP, AFJ, RJS, MS, SG, AR-T, AHM
Formal analysis: DDS, MVFR, AHM
Visualization: DDS, AHM
Funding acquisition: AHM
Writing – original draft: DDS, AHM
Writing – review & editing: DDS, MVFR, SG, AR-T, AHM

## Competing interests

Authors declare that they have no competing interests.

## Data availability

All sequence data generated in this study have been deposited to the National Center for Biotechnology Information Sequence Read Archive under accessions BioProject ID PRJNA870104.

## Supplementary Materials

Materials and Methods
Supplementary Results
Supplementary Code
Supplementary Figures S1 to S8
Supplementary Tables S1 to S5
Supplementary References (*36–44*)

### Materials and Methods

#### Ethical statement

All procedures conformed to guidelines established by the U.S. National Institutes of Health and have been approved by the Cornell University Institutional Animal Care and Use committee (IACUC: Protocol #2015-0060). The Institutional Animal Care and Use Committee at the University of Nebraska-Lincoln approved all procedures involving ex-germ-free mice (Protocol 1700).

#### Animal Husbandry

Donor fecal samples were collected from five lineages of *Mus musculus domesticus* (NY2, NY4, FL1, LEWES/EiJ, and C57BL/6J), two lineages of *Mus spicilegus* (ZBN and SPI/TUA), one strain of *Mus pahari* (PAHARI/EiJ), and one lineage of *Peromyscus maniculatus* (referred to here as PMAN). All *Mus* lines were maintained by sibling mating in a common laboratory environment using standard mouse husbandry procedures for at least 20 generations. Wild-derived inbred *M. m. domesticus* lines NY2 and NY4 were derived from distinct initial pairings of wild mice trapped from Saratoga Springs, NY, USA, whereas the FL1 line was derived from a pair of wild mice trapped in Gainsville, FL, USA. Additional house mouse strains of *M. m. domesticus* (LEWES/EiJ and C57BL/6J) as well as *M. spicilegus* (ZBN and SPI/TUA), and *M. pahari* (PAHARI/EiJ) lines were originally obtained from The Jackson Laboratory (Bar Harbor, Maine, USA), where the lines were maintained without rederivation. These lines from Jackson Laboratory were subsequently maintained at Cornell University alongside the *domesticus* lines for >10 generations. The *Peromyscus maniculatus* line was originally derived by the Peromyscus Genetic Stock Center (University of South Carolina, Columbia, SC, USA). All mice were fed standard lab-mouse chow (~19% Crude Protein, ~6% fat, 44% carbohydrate, ~18% fiber). Fecal samples from individual mice were weighed and combined into mixtures at Cornell University, then shipped on dry ice overnight to the University of Nebraska-Lincoln. Our previous work has shown that >85% of the family-level taxonomic diversity in mouse gut microbiota remains viable after freezing (i.e., >85% of the bacterial families detected in fecal samples we cultivated under anaerobic conditions on a panel of 9 media) (*35*). Fecal slurries were suspended in reduced, sterile PBS and gavaged into individual germ-free mice.

Germ-free recipient *M. m. domesticus* (C57BL/6J and *Rag1*^−/−^) were born and reared in flexible film isolators and maintained under gnotobiotic conditions (temperature 20 °C, relative humidity 60%, 14 h light/10 h dark cycle) by the Nebraska Gnotobiotic Mouse Program at the University of Nebraska-Lincoln. For experiment 1 (*i.e*., experiment shown in Fig. 1C), male and female germ-free mice were transferred from isolators to sterile, individually-ventilated cages with high-performance filter lids at the time of colonization. For experiment 2 (*i.e*., experiment shown in Fig. 2A), germ-free mice were colonized and maintained in cages within gnotobiotic isolators for the duration of the study. All mice were fed autoclaved chow (LabDiet 5K67, Purina Foods, St. Louis, MO, USA) *ad libitum*.

#### 16S rRNA Amplicon Profiling

DNA was extracted from all samples using the DNAeasy PowerLyzer PowerSoil Kit from QIAGEN (Hilden, Germany). Donor fecal samples, inoculum mixtures, and output fecal samples from ex-germ-free mice used in experiment 1 were sent to the Microbiome Core Lab located at the Jill Roberts Institute for Research in Inflammatory Bowel Disease of Weill Cornell Medicine (New York, NY, USA) for sequencing. Briefly, 16S rRNA (V4-V5) amplicon libraries were prepared using 515F-926R primers developed by the Earth Microbiome Project (*36*).

Library pools were then sequenced in one PE250 run on an Illumina MiSeq (San Diego, CA, USA).

DNA from donor fecal samples and output fecal samples from ex-germ-free mice used in experiment 2 were sent to the Integrated Microbiome Resource at Dalhousie University (Halifax NS, Canada). 16S rRNA (V4-V5) amplicon libraries were prepared using 515F-926R primers. Libraries were sequenced in a single PE300 Illumina MiSeq run with a V3 reagent chemistry (San Diego, CA, USA).

Raw reads from both experiments were denoised into ASVs using the DADA2 pipeline (v1.14.0) (*37*). Following generation of an ASV table, sequences were chimera-checked, and those remaining that were not 230–235 bps in length were removed. Taxonomic assignments were made using the RDP classifier and both GreenGene database (13_8) (*38*) and the SILVA nr database (v132) (*39*) (Table S5). ASVs that were not identified as Domain Bacteria or Domain Archaea were excluded from further analyses. A phylogenetic tree of the total set of ASVs was inferred using the fragment insertion function (*40, 41*) in QIIME2 (v2019.1) (*42*) and the full GreenGenes 13_8 tree. A mapping file, the ASV table, the taxonomy table, and the phylogenetic tree were imported into R and combined into a single phyloseq object (v1.28.0) (*43*).

#### 16S rRNA Gene Copy Number Quantification

The Femto Bacterial DNA Quantification Kit (Zymo, Irvine, CA, USA) was used to quantify the bacterial load of donor and ex-germ-free mouse fecal samples on the Applied Biosystems QuantStudio 7 Pro Real-Time PCR System (Waltham, Massachusetts, USA). Extracted DNA was diluted and quantified using the standard analysis protocol. Technical replicates (*i.e*., replicates of the same DNA sample) were averaged.

#### Data Analysis

Shannon Diversity index and the Jaccard similarity index were calculated performed using the phyloseq R package. Pielou’s Evenness index was calculated using the microbiome R package (v1.16.0). Additional statistical analyses including Chi-Squared, Fisher’s Exact, Exact Binomial and Wilcoxon tests were performed in R. PERMANOVA tests were performed using the ‘adonis’ function in the vegan R package (v2.5.5). PCoA ordinations, boxplots, and scatter plots, were generated using the ggplot2 package in R (v3.2.0). Phylogenetic trees were created using the iTOL online tool (v6) (*44*).

#### Local Adaptation Index Calculations

To assess the degree of local adaptation of microbiota to house mice, we defined the Local Adaptation Index (LAI) as the Jaccard similarity of the microbiota of the ex-germ-free mouse to that of the native house-mouse donor minus the similarity of the microbiota of the ex-germ-free mouse to that of the non-native donor.

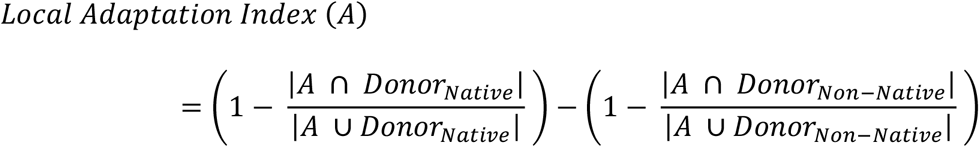

Larger positive LAI values indicate that a given profile from ex-germ-free mice was more similar to its native donor than it was to its non-native donor, indicative of local adaptation of the gut microbiota to the *M. m. domesticus* gut.

The Local Adaptation Index was also calculated for each microbiota simulated under a neutral model of community assembly. These values were used as the expected LAI values for ex-germ-free mice given neutral assembly. Therefore, the difference between the observed LAI for an ex-germ-free mouse fecal microbiota and the expected LAI for the microbiota (*i.e*., Observed LAI – Expected LAI) provided a test statistic for the degree of competitive advantage of the native microbiota over the non-native microbiota.

#### Simulation of Expected Composition Under Neutral Assembly

We simulated the microbiota composition expected under a model of neutral community assembly for each ex-germ-free mouse inoculated with a mixture of native and non-native microbiotas. For these analyses, we incorporated both the compositions of the donor samples estimated from 16S rRNA gene sequencing and the bacterial loads of the donor samples estimated from qPCR of the 16S rRNA gene. We measured the bacterial load of each donor sample using qPCR and multiplied the copies per gram by the mass of each fecal pellet included in the inocula mixture to yield an estimate of the total number of 16S rRNA gene copies in the mixture from each donor. The fractional proportion of each donor’s microbial load was then used to weight the subsampling of the microbiotas from each donor strain in order to generate the expected microbiota composition in the ex-germ-free mouse under neutral assembly. For analyses of selection on individual ASVs, the expected relative abundance of each ASV for each mixture was calculated based on neutral assembly using the weightings from the observed 16S rRNA gene profiles and loads in the donor fecal samples. Further details of these analyses, as well as a fully reproducible workflow, are available as a Supplementary Code file.

#### SourceTracker Analysis

SourceTracker (v1.0.1) (*29*), a Bayesian method for estimating the likelihood of microbes within a sample having originated from each of multiple sources, was used to assess the most likely origin of ASVs in each fecal sample from ex-germ-free mice. Here we estimated the most likely origin of ASVs in every fecal pellet from ex-germ-free mice using the corresponding native and non-native donors as the “source” communities. Sample and donor communities were rarefied to the same, minimum sequencing depth prior to SourceTracking to control for differences in sequencing depth between donors. SourceTracker was employed with default settings.

#### Code availability

A detailed description of these analyses, along with all datasets and analysis code, is available at github (https://github.com/DanielSprockett/CU01_Local_Adaptation/) and in Supplementary Materials.

---

**Fig. S1.**
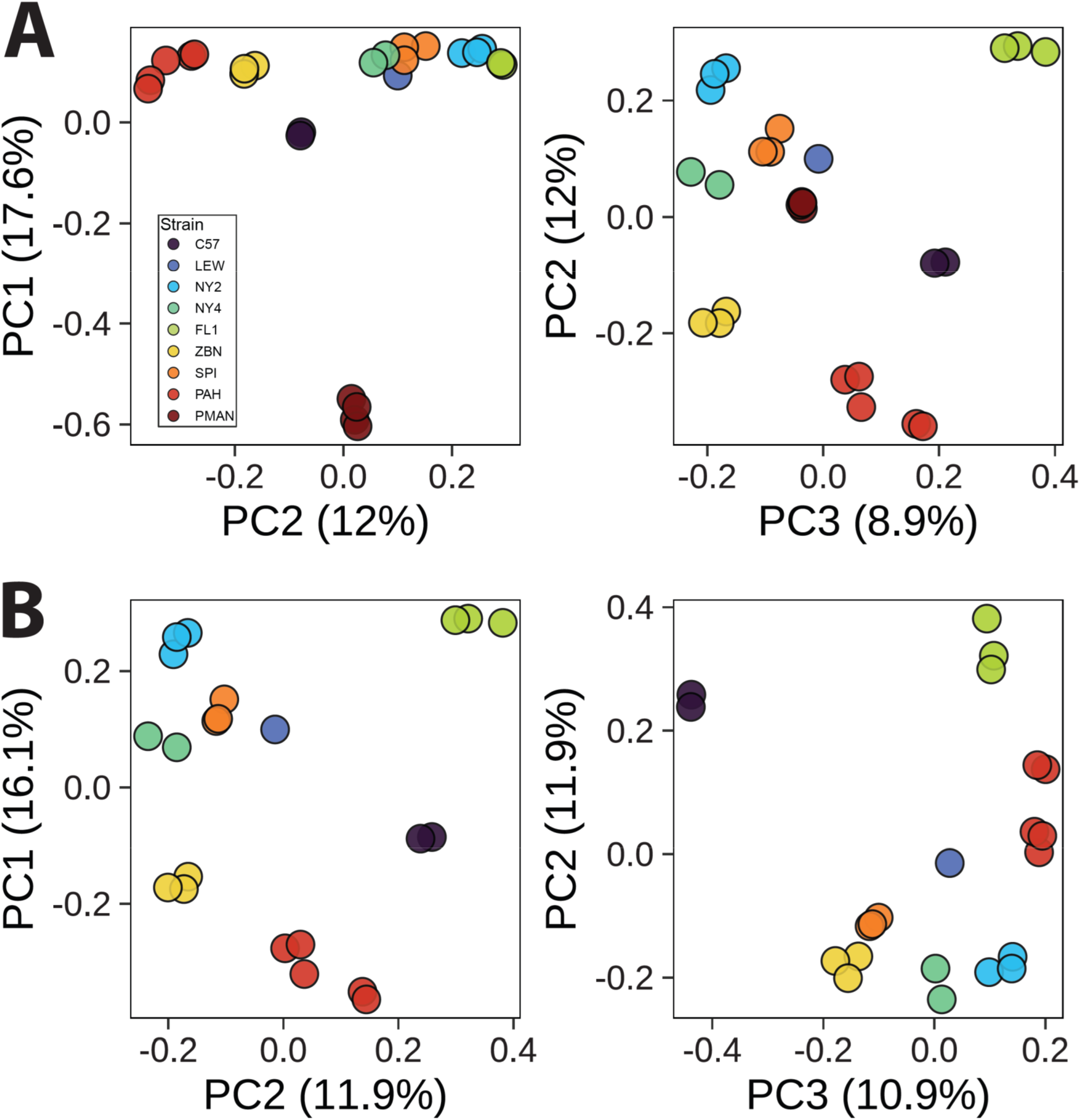
Microbiotas from diverse mouse lineages were compositionally distinct. (**A, B**) Principal Coordinate Analysis plots show similarity of microbiotas among wild-derived mouse lineages based on the Jaccard similarity index of 16S rRNA amplicon profiles generated from donor fecal samples (i.e., fecal samples gavaged into germ-free mice). Plots show all donor mouse samples used in this study (**A**) or only donor samples from *Mus* (**B**). Colors indicated the rodent strain from which the fecal sample originated. PERMANOVA p <0.001 for A and B.

**Fig. S2.**
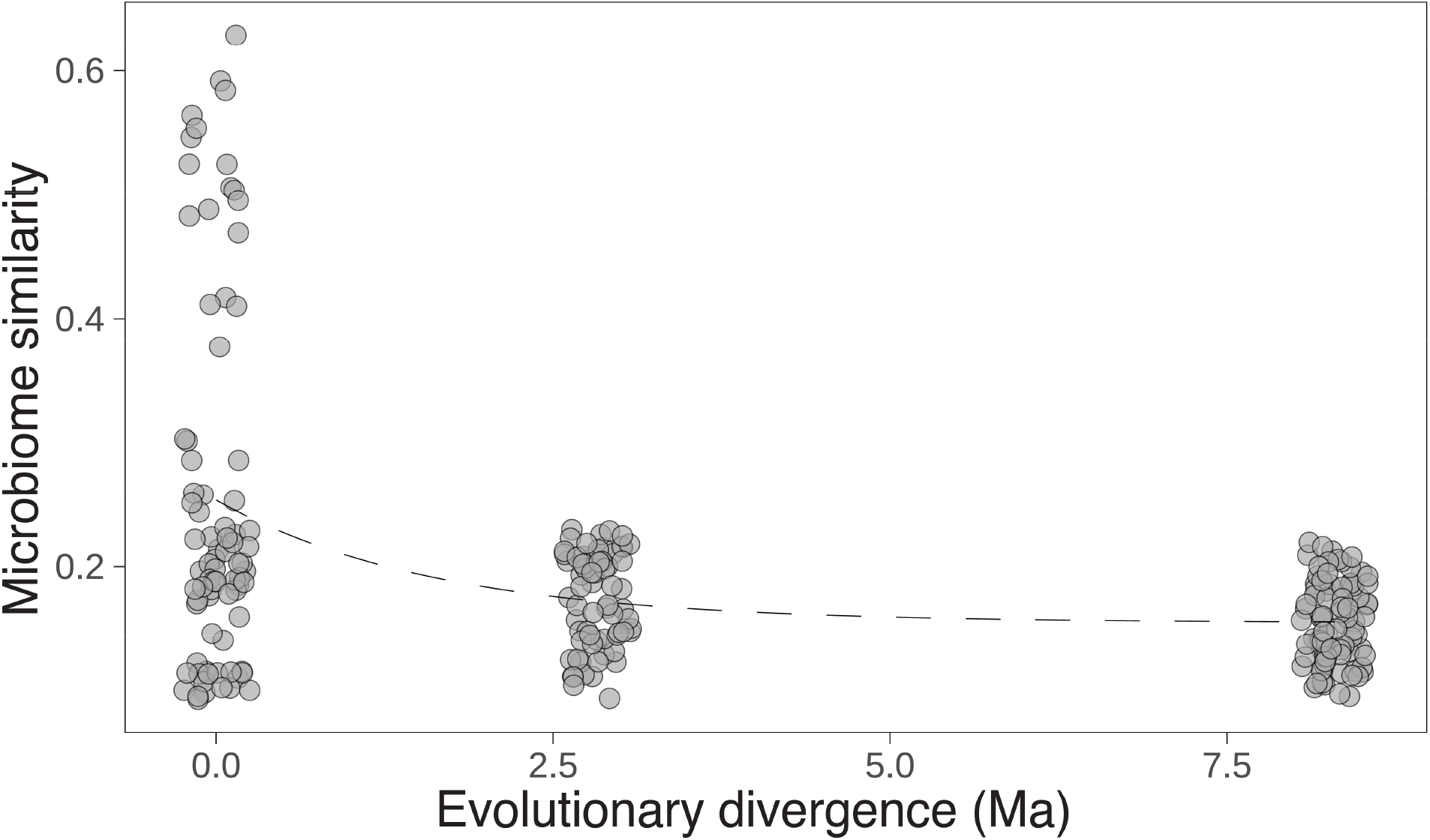
Microbiotas from *Mus* lines recapitulated donor phylogeny. Scatterplot shows the negative relationship between microbiota similarity and host evolutionary divergence time (Ma, millions of years ago) among different mouse lines in the genus *Mus* based on the Jaccard similarity index. Dashed line indicates exponential decay (p <0.001, R^2^ = 0.19). Boxplots in inset show Jaccard similarities between pairs of samples from the same host strain (leftmost), different host strains from the same species (middle), and different *Mus* species (right). For each boxplot, the center line denotes the median, and the lower and upper hinges correspond to the first and third quartiles, respectively. Wilcoxon tests, FDR-adjusted p-values **** <0.0001.

**Fig. S3.**
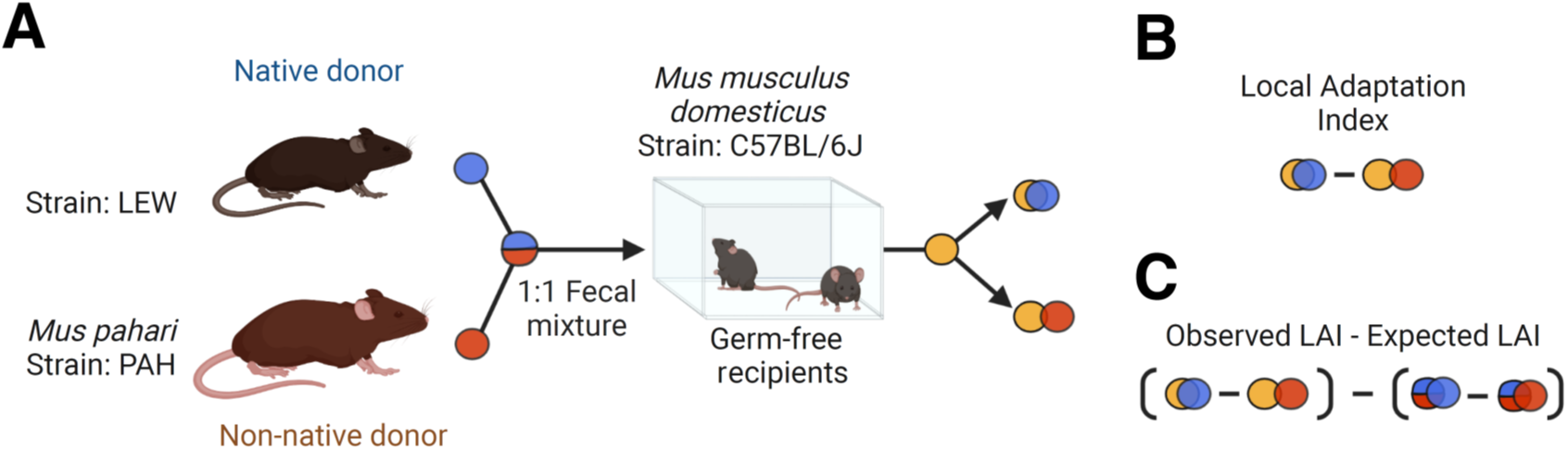
Calculation of Local Adaptation Index. (**A**) Fecal pellets from native and non-native donors were mixed 1:1 by weight and then gavaged into germ free mice. (**B**) Microbiota similarities between the microbiotas of ex-germ-free mice and donors were calculated. The Local Adaptation Index of a microbiota of an ex-germ-free mouse was defined as the dissimilarity of the microbiota to the native donor minus the dissimilarity of the microbiota to the non-native donor. (**C**) Calculating the difference between the observed LAI and the expected LAI under neutrality (i.e., even mixtures of native and non-native microbiota, accounting for variation in microbial load and weight of donor fecal pellets) provided a test statistic for local adaptation.

**Fig. S4.**
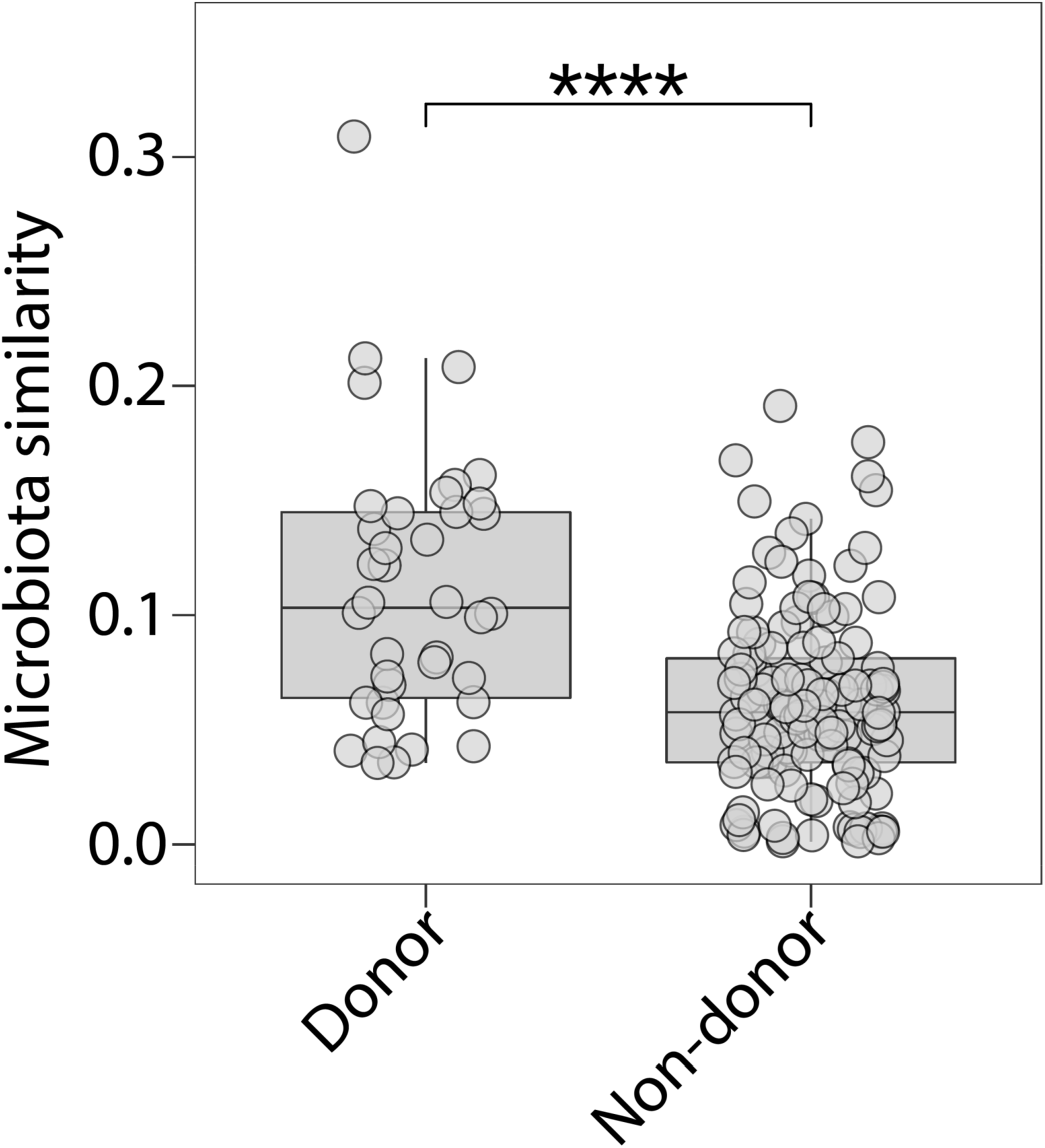
Microbiota similarity of ex-germ-free mice to donor mouse fecal samples. Boxplots show, for experiment 1 (Fig. 1), the Jaccard similarity between ex-germ free mice and their corresponding donors as well as the Jaccard similarity between ex-germ free mice and other donors. Significance of Wilcoxon tests is indicated by asterisk; **** *p*-value <0.0001.

**Fig. S5.**
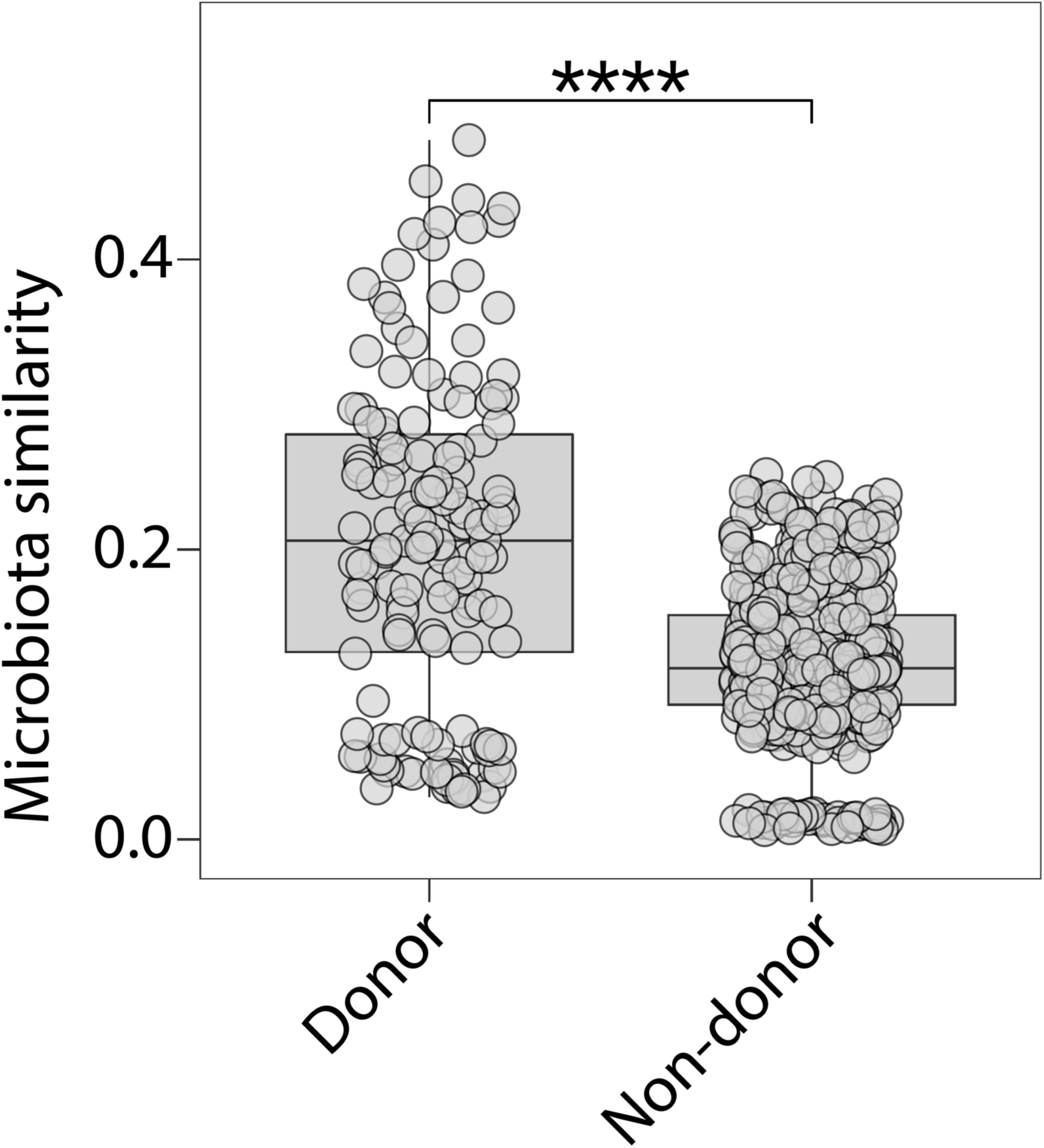
Microbiota similarity of ex-germ-free mice to donor mouse fecal samples. Boxplots show, for experiment 2 (Fig. 2), the Jaccard similarity between ex-germ free mice and their corresponding donors as well as the Jaccard similarity between ex-germ free mice and other donors. Significance of Wilcoxon tests is indicated by asterisk; **** *p*-value <0.0001.

**Fig. S6.**
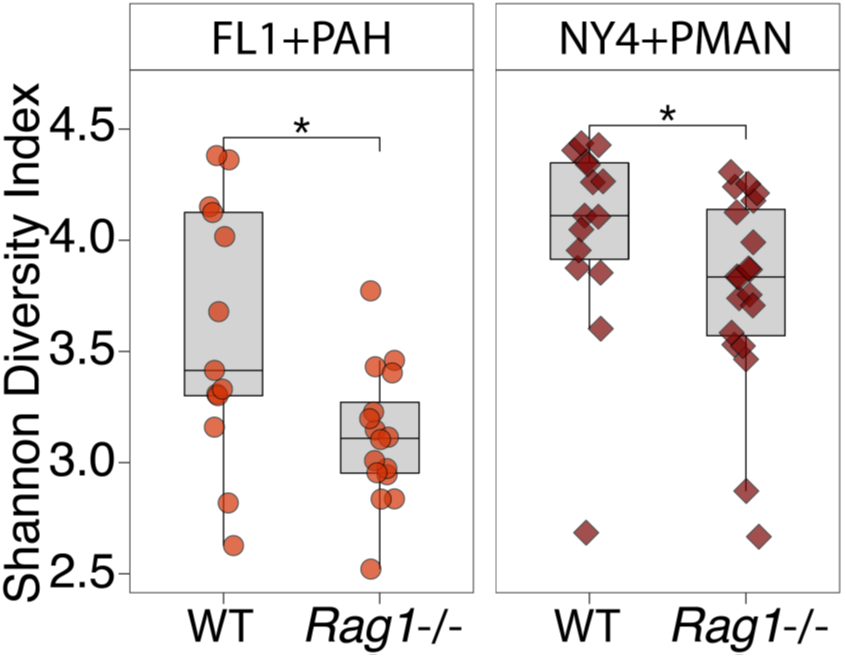
Alpha diversity differs between WT and *Rag1*^−/−^. Boxplots show, for experiment 2 (Fig. 2), the alpha diversity (Shannon) of ex-germ free mice. Significance of Wilcoxon tests is indicated by asterisk; **** *p*-value <0.0001.

**Fig. S7.**
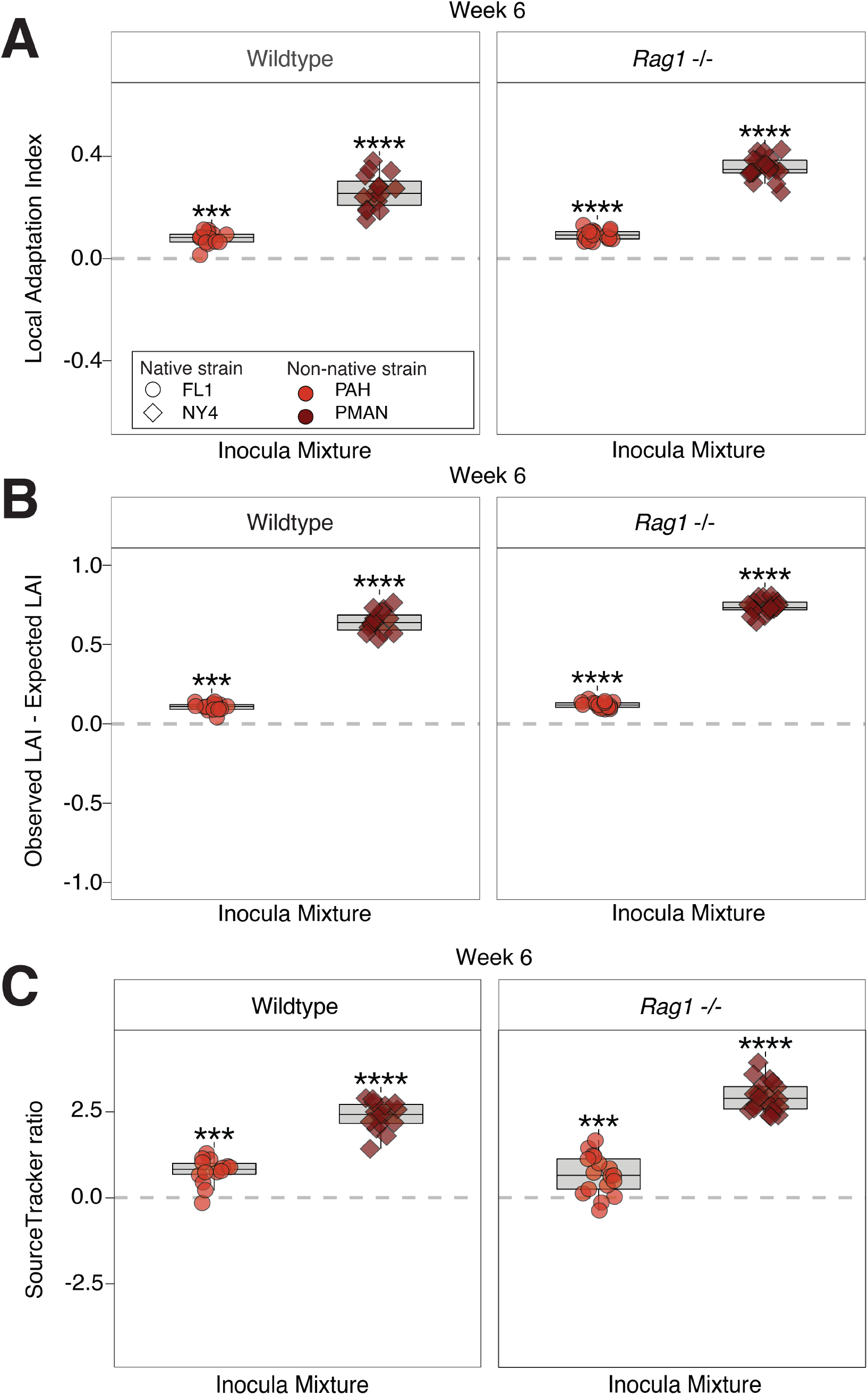
Adaptive advantages of native over non-native microbiotas persisted to week 6 in both WT and *Rag1*^−/−^ mice. (**A**) Boxplots show positive Local Adaptation Index (LAI) values, indicating that microbiotas of ex-germ-free WT and *Rag1*^−/−^ mice at week 6 were more similar to the microbiotas of their native donors than they were to those of their non-native donors. (**B**) Boxplots show positive differences between observed LAI values and LAI values expected under neutral assembly. (**C**) Boxplots show positive log_10_ transformed ratios between the percentage of the microbiota derived from the native donor (estimated with SourceTracker) to the percentage derived from the non-native donor. Shapes and colors denote identities of native and non-native donors, respectively. Lines connect samples from the same mouse. Wilcoxon test for non-zero mean, FDR-adjusted p-values * <0.05, ** <0.01, *** <0.001, **** <0.0001. For each boxplot in (**A**–**C**), the center line denotes median, and lower and upper hinges correspond to first and third quartiles, respectively.

**Figure S8.**
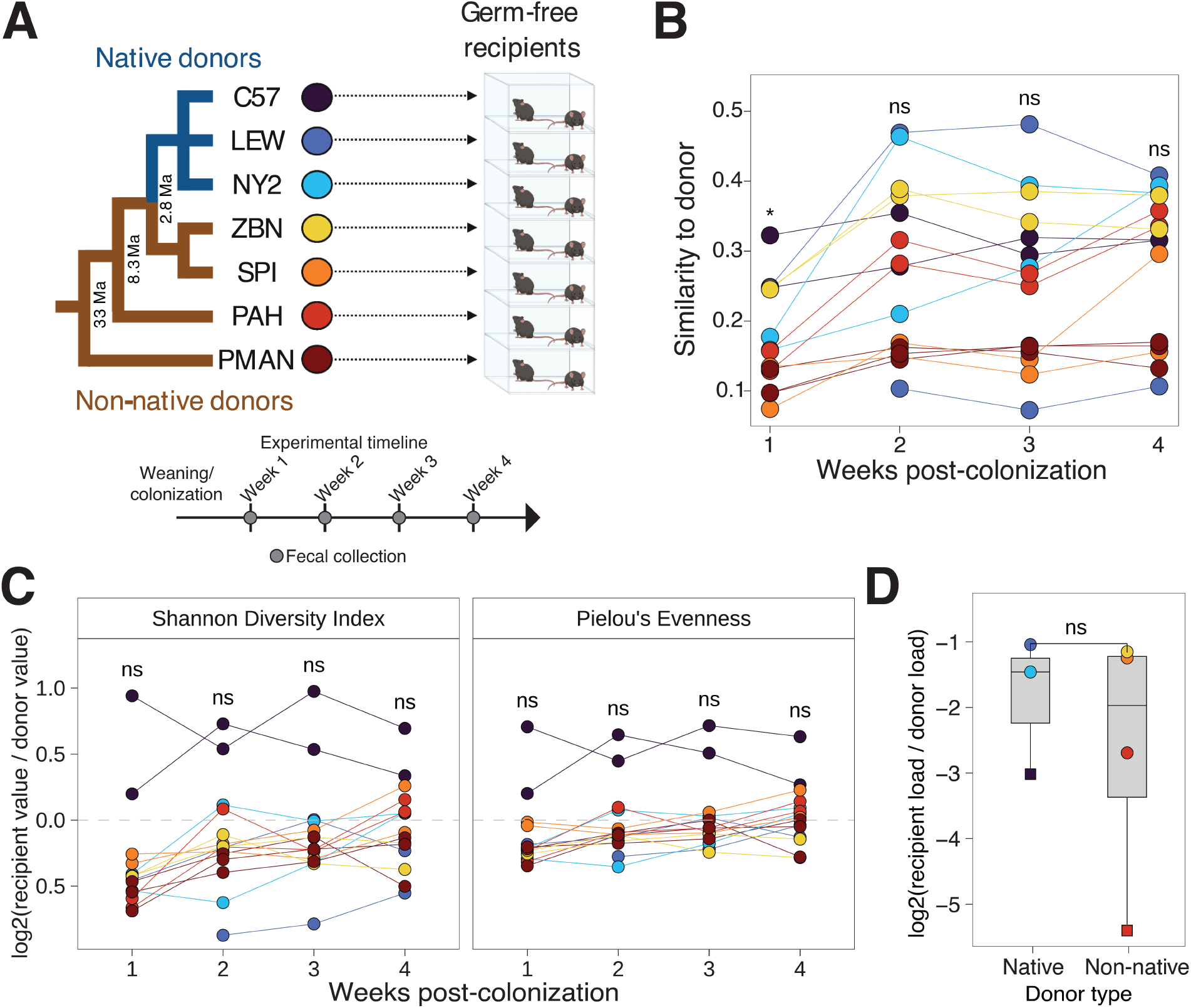
Native and non-native microbiotas colonized germ-free mice to comparable degrees when inoculated individually. (**A**) Cartoons show experimental design. Fecal pellets from three native *Mus musculus domesticus* lines and four non-native mouse strains were inoculated directly into germ-free mice. Fecal pellets were collected from ex-germ-free mice weekly for 4 weeks. (**B**) Lines show Jaccard distanced of microbiotas from ex-germ-free mice to their donor microbiotas over the 4-week experiment. Each line corresponds to an ex-germ-free mouse, and each point the compositionally dissimilarity between the microbiota from an exgerm-free mouse and the donor microbiota that the mouse received. (**C**) Lines show log_2_-transformed ratios of Shannon’s Diversity Index and Pielou’s Evenness between the microbiotas of ex-germ-free mice and their donor microbiotas during the 4-week experiment. (**D**) Boxplots show log_2_-transformed ratio between the bacterial loads per gram of feces in ex-germ-free recipients and donors. Ex-germ-free mice that received house-mouse microbiota are plotted separately from those that received non-native microbiotas. For panels (**A**–**D**), colors denote the identities of the mouse donors. Lines connect samples from the same mouse. Wilcoxon test, FDR-adjusted p-values ns = not significant, ns = not significant, * <0.05.

Table S1. Metadata for rodent donor and recipient samples.

Table S2. Local Adaptation Index values from experiment 1.

Table S3. Bacterial load estimates using 16S rRNA gene targeted qPCR.

Table S4. Local Adaptation Index values from experiment 2.

Table S5. Selection results and taxonomy assignments for ASVs.

## Notes

https://github.com/DanielSprockett/CU01_Local_Adaptation

https://zenodo.org/record/7003965#.YwO2fezML0o

### Competing Interest Statement

The authors have declared no competing interest.

